# Machine learning assisted multi-level differentiation of blood cells using multi-frequency electro-mechanical impedance cytometry

**DOI:** 10.64898/2026.11.12.766153

**Authors:** Ishita Bansal, Suraj K Suresh, Nitish Sagar, Prosenjit Sen

## Abstract

We report a multi-frequency microfluidic impedance cytometer with multimodal measurement of electrical and mechanical cell properties. Using the large feature set from the multimodality, we implemented machine learning assisted differentiation of cells. We have demonstrated a constriction-equipped channel to study the mechanical cell properties, and integrated multiple microelectrodes that capture electric field information pertaining to various stages of cell transit. Our system works over a wide frequency range from 800 kHz to 8 MHz.

We developed a machine learning workflow to classify cells using our multi-modal data. Using this platform to show blood cell differentiation, our system shows minimum F1 score to be 0.92, minimizing both false negatives and false positives, which is important for medical diagnosis. Furthermore, we also identified the presence of two populations in the mixed sample of neutrophils and PBMCs. Our workflow allows classification of cells in a new sample along with identification of top 5 distinguishing parameters out of 70+ features extracted from the raw impedance signal.

## 1. Introduction

Impedance cytometry has emerged as a rapid label-free technique to conduct cell and particle studies. Began as early as 1950s through Coulter principle for counting cells using electrical impedance measurement with a direct current (DC) signal, the field of impedance cytometry has expanded to use alternating current (AC) signal at multiple frequencies to probe various parts of the cell. Such impedance cytometers are now integrated into microfluidic devices to enhance throughput and portability while examining the biophysical properties of cells at a single-cell level. Electrical and mechanical properties of cells are widely studied label-free biophysical markers for understanding the physiological status of cells.^[1]^

Leveraging high-throughput capability, non-invasive nature and label-free processing, microfluidic impedance cytometers are actively used to differentiate between multiple kinds of cells using their electrical properties. Tang et al.^[2]^ analysed various dielectric parameters such as cell diameter, impedance amplitude and phase shift at multiple frequencies to distinguish between white blood cells and four kinds of tumor cells (A549, H460, MCF7, and H226). Zhu et al.^[3]^ detected circulating tumor cells in peripheral blood by a liquid biopsy instrument employing inertial microfluidics and impedance measurement using facing electrodes. Other studies have been conducted to distinguish between cells such as live vs dead yeast^[4]^, identification of algae and cyanobacteria in freshwater to employ appropriate control strategies ^[5]^, and identification and differentiation between two kinds of parasitic protozoa in water bodies^[6]^ using impedance measurement. However, electrical properties alone often fail to provide a unique “fingerprint” for specific cell types, leading to overlapping distributions.

Another biophysical property of interest to engineers is the mechanical property of cells. The mechanical properties of cells such as elasticity and stiffness^[7]^ provide information about the response of cells to stress. Conventionally, micropipette aspiration, atomic force microscopy, optical tweezers and magnetic tweezers have been used to measure the stiffness of cells. However, these techniques suffer from low throughput (10-20 cells/hour)^[8], [9], [10]^ and pose a risk to cell viability. Deformability cytometry, utilising microfluidic constriction channels, measures stiffness of cells in flow at high throughput (∼100-1000 cells/sec).^[11]^ Constriction-based microchannel design deforms the cells as they pass through a smaller-than-cell constriction, leading to different transit times for varying stiffnesses of cells. These transit times are measured using high speed imaging. The mechanical properties of red blood cells^[12], [13], [14]^, cancer cells^[15], [16]^, and white blood cells ^[17]^ have been studied using deformability cytometers. Recent approaches such as real-time deformability cytometry (RT-DC) enabled high-throughput (∼1000 cells/sec)^[11]^ mechanical phenotyping of cells using high-speed imaging and real-time image analysis. However, the primary difficulty with such deformability measurements is the high cost and complexity of imaging setups and time-consuming image processing algorithms.

Integrating electrical and mechanical single-cell measurements is essential to improve biophysical phenotyping. Having multiple modalities of measurement expands the feature dimension for effective machine learning-based cell differentiation. Recently, a shear-flow deformability cytometer combined impedance sensing with flow-induced deformation at throughputs of ∼100 cells/sec^[18]^. However, they employed complex multi-step fabrication process for facing electrodes which requires precise alignment. Cui et al. ^[19]^ developed a device to automatically trap cells, deform them and measure their impedance, allowing concurrent measurement of mechanical and electrical properties of the cells, but lacked high throughput (∼10 cells/s). Zou et al. ^[20]^ demonstrated an electro-optical deformability cytometer that tracks the changes in cell shapes electrically and optically during shear deformation. However, shear deformation does not properly squeeze cells and may not fully probe cytoskeletal or membrane mechanical limits. Zhou et al. ^[21]^ demonstrated a combined electrical-mechanical cytometer by deploying two sets of electrodes, one at the entrance and other at the exit of the constriction, to account for the entry and transit times of cells. However, their technique heavily depends on passage time, which gets affected by flow rate fluctuations, to understand mechanical properties and fails to decouple cell stiffness and cell size.

To successfully classify cells using a machine learning workflow, a large number of distinct cell features are required. Integrating constriction with electrodes allows us to capture both the electrical and mechanical properties of cells during the different stages of cell transit. We demonstrate a high-throughput multi-frequency impedance cytometer that allows us to capture maximum number of distinct features related to both the electrical and the mechanical cell properties. We present a unique coplanar electrode layout with four sets of electrodes strategically placed around the constriction, which allow simultaneous multifrequency electrical measurements in the native, deformed and post-deformed states of cells. The presence of two pairs of electrodes before the constriction allows the measurement of cell velocity in its undeformed state which is essential for long experiments where flow rate fluctuates during the experiment. We have also placed the constriction electrode at the centre of the constriction to measure cell impedance in its maximum stress condition. We believe that this distinctive design will enhance our understanding of internal cellular structures and enable reliable and high-throughput single-cell electro-mechanical analysis.

## 2. Device concept

**Figure 1(a)** shows the schematic illustration of the system. Figure 1(b) shows the device configuration. There are two sets (A and B) of electrodes with each set consisting of two pairs. The first set (A1 and A2) is present before the constriction measuring two main parameters: (i) the electrical properties of cells in their native state; and (ii) the velocity of cells flowing through the straight channel. The experiment is conducted at three distinct frequencies: (a) cell volume at 800 kHz, (b) cell membrane at 2.3 MHz, and (c) cytoplasm at 8 MHz ^[22]^. The parameters of interest from the first electrode set are:

1. amplitude at before constriction electrodes, Amp BC+ and Amp BC-, taken as the impedance magnitude at A1 and A2; and
2. passage time, T_p_, measured as centre-to-centre time as the cell passes from A1 to A2.

**Figure 1:**
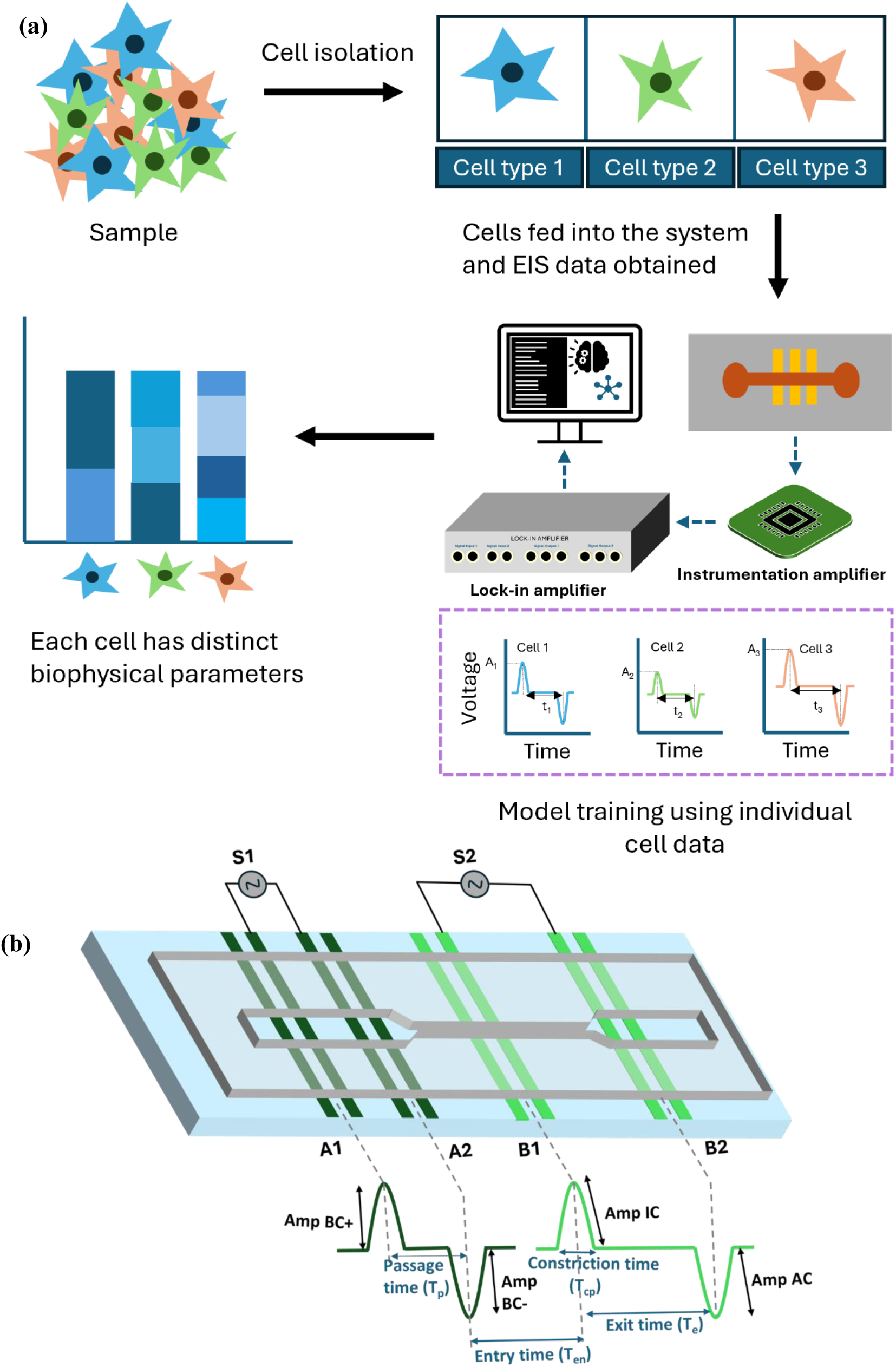
(a) Schematic illustration of the working of the system: A sample of cells comprising of multiple cell types is taken and individual cell types are isolated and collected separately, suspended in buffer. Individually, each cell solution is ffown through the device, and the electrical impedance is collected. Using the impedance data from all the cell types, a machine learning model is trained which is then used to classify cells. (b) Device design with electrical impedance signatures corresponding to each pair of electrodes.

We derived another parameter, Max Amp BC, which is calculated as the maximum value of amplitude from BC+ and BC-electrode at 800 kHz. This was done to consider the value of amplitude when the cell is closest to the electrodes, thereby giving a clear picture of the cell volume. We used the cube root of Max Amp BC, i.e., Max Amp BC^1/3^, as the cell diameter equivalent.

The second set of electrodes (B1 and B2) consist of one pair of electrodes (B1) placed inside the constriction and the second pair (B2) just after the constriction. The benefit of such a placement is three-fold: (i) B1 measures the electrical property of the cell in its deformed state, (ii) B2 measures the property of the cell post deformation, and (iii) both B1 and B2, in conjunction, measure the time it takes for the cell to exit the constriction. The parameters extracted from the second set of electrodes are:

1. entry time, T_en_, measured from centre-to-centre of A2 and B1, as the cell passes from A2 to B1 entering the constriction.
2. amplitude at inside constriction electrodes, Amp IC, taken as the impedance magnitude at B1.
3. amplitude at after constriction electrodes, Amp AC, taken as the impedance magnitude at B2.
4. constriction passage time, T_cp_, taken as the time it takes for the cell to transit through the B1 electrode pair; and
5. exit time, T_e_, measured from centre-to-centre of B1 to B2, as the cell passes from B1 to B2 exiting the constriction.

Furthermore, the above parameters were used to generate other parameters such as membrane opacity and cytoplasmic opacity, measured as the ratio of amplitude at 2.3 MHz to 800 kHz and 8 MHz to 800 kHz respectively. For machine learning-based analysis, the electrode placement is selected such that we obtain the maximum number of independent features as the cell traverses through the microfluidic channel with constriction.

## 3. Results and Discussion

We conducted the study in three levels: the first level was differentiating between the three major blood cells, namely the neutrophils, PBMCs and the RBCs flowed sequentially; the second level was differentiating between neutrophils and PBMCs from their 1:1 mixed population; and the third level was further differentiation of the sub-populations of PBMCs. In the first study, neutrophils were flowed at the beginning, considering their short lifespan outside the human body at room temperature and pH, followed by PBMCs and lastly, RBCs. The device used for all these three studies had the dimension of 20 um (width) x 15 um (depth) straight channel and 7 um (width) x 15 um (depth) constriction.

### 3.1 Sequential flow of Neutrophils, PBMCs and RBCs

Neutrophils and PBMCs were extracted from blood using the density gradient centrifugation method and diluted in PBS buffer. RBCs were obtained by diluting the blood in PBS buffer. The final cell concentration was kept at 1.5×10^3^ cells/µL. The cells were introduced in the device using Chemyx syringe pump at a constant flow rate of 1 µL/min. The experiment was conducted by initially flowing neutrophils in the device, followed by PBMCs and lastly RBCs, each cell type flown for 30 minutes and peak data obtained from HF2LI and saved into the computer. The analysis was done offline after the experiment. Equal number of all the three cell types were used for machine learning classification.

#### 3.1.1 Raw peak analysis

Figure 2 shows the process of analysis of peaks obtained from the lock-in amplifier. The raw peaks were first baseline corrected using a moving average filter. Next, the peaks were identified individually, both positive and negative, using Python’s SciPy library functions. The individual peaks were then paired with each other using distance and amplitude constraints, followed by peak-to-peak pairing of before constriction (BC) peaks and inside-and-after constriction (IAC) peaks using similar constraints. Lastly, the peaks were fitted with gaussian functions to extract transit times and peak widths. Figure 2(b) shows the R^2^ box plot of the gaussian fitting of all the four kinds of peaks: positive peak 1 (BC+), negative peak 1 (BC-), positive peak 2 (IC) and negative peak 2 (AC): with a high mean R^2^ value (R^2^ >0.91) showcasing a good fit.

**Figure 2:**
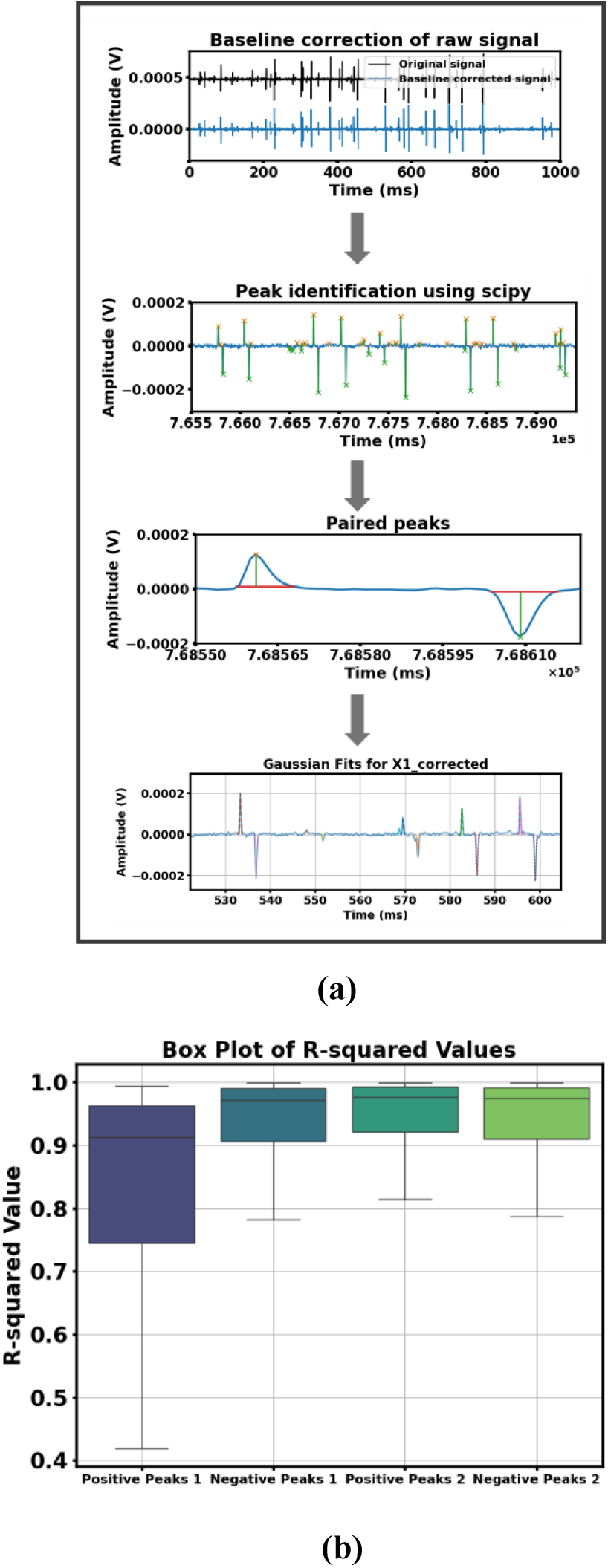
(a) Basic blocks of the peak detection algorithm, starting with baseline correction using a moving window average, then identifying positive and negative peaks using SciPy library functions, followed by peak pairing using amplitude and distance constraints and later, gaussian fitting of individual peaks to extract times and peak widths; (b) box plot of R^2^ values for each of the gaussian fit

#### 3.1.2 Machine learning analysis

After the preliminary analysis, we created a machine learning workflow for understanding the importance of various features towards cell differentiation and validating our technique to unseen data. Figure S4 shows the ML workflow developed for advanced analysis. The input to the algorithm was a 72-column concatenated feature DataFrame of all three cell types, obtained as the output from the peak analysis. The heatmap of the original 72 features over a few hundred cells is shown in figure S4(a). Beginning with an unsupervised learning approach to identify the presence of different clusters, we used this vast feature space as input to a dimensionality reduction technique: uniform manifold approximation and projection (UMAP) which revealed three distinct clusters.

UMAP projects a high dimensional data into a 2D or a 3D space, effectively reducing the dimension of the data and allowing for better visualisation of the data. It works by connecting individual data points to their neighbours using a fuzzy topological structure, where the connection strength represents the probability of the two points being connected. UMAP then optimises the representation of the data in the 2D or 3D space.^[23]^ Since the data generated by our device is multi-dimensional, using a dimensionality reduction technique was essential in finding clusters in the data and visualise them. Compared to other similar techniques such as t-distributed stochastic neighbour embedding (t-SNE), UMAP is significantly faster and preserves the global structure of the data. UMAP is also better in preserving the shape of the data than principal component analysis (PCA) as PCA has a linear approach, but UMAP has a non-linear approach to dimensionality reduction which helps it to be used on complex data.

Figure 3(a) shows the 2D density scatter map colour coded with the cell type label. UMAP1 and UMAP2 are two dimensions generated from the 72 dimensions(features) fed as input to the UMAP algorithm. The presence of three clusters in the figure confirms the presence of three distinct cell populations in our data. We gathered the top features contributing to each of the two UMAP coordinates by running a random forest regressor trained on the UMAP coordinates as target labels. Random forest is a supervised machine learning algorithm comprising of an ensemble of decision trees which together vote for the most popular class.^[24]^ Since UMAP is a non-linear algorithm, a non-linear regression algorithm such as random forest was selected to extract the features most responsible for the UMAP coordinates. Figure 3(b) shows the bar graph for the top 15 drivers of the UMAP1 and UMAP2 coordinates across their relative importance obtained through Gini importance. Gini importance, also known as Mean Decrease in Impurity, quantifies the contribution of each feature to the prediction power of a model by measuring the total reduction in impurity brought about by that feature. Y Amp BC-2.3 MHz and T_e_ were found to be the topmost drivers for UMAP1 and UMAP2 respectively. The impact of the two drivers on the UMAP plot is depicted in figure 3(c).

**Figure 3:**
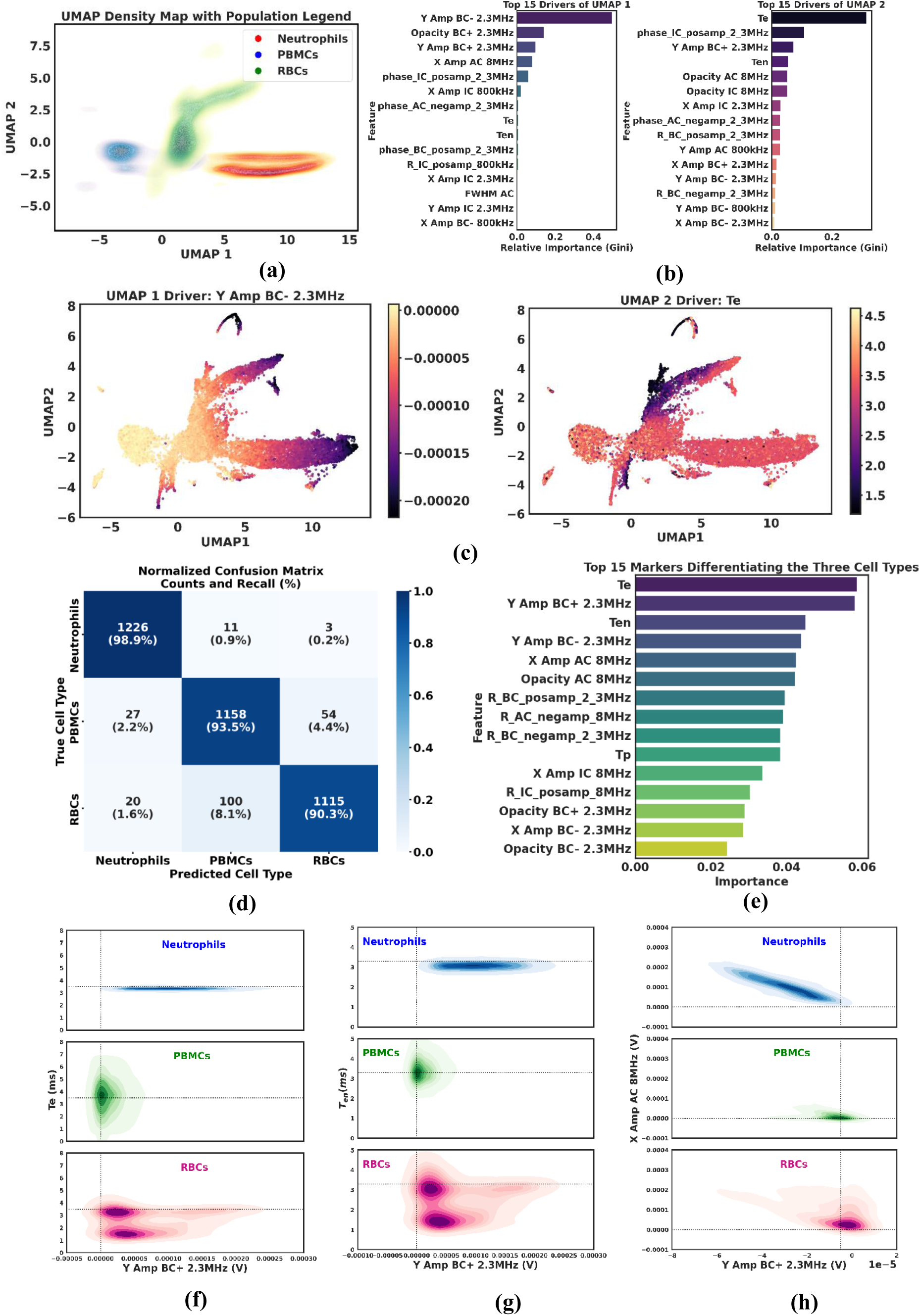
ML analysis results for Neutrophils, PBMCs and RBCs (a) UMAP density map showing the presence of three distinct clusters; (b) top 15 features contributing to each of the UMAP coordinates obtained using Gini importance; (c) Overlap of the top drivers of UMAP1 and UMAP2 on the UMAP projection plot showing the variation of the driver values throughout the UMAP projection space; (d) confusion matrix for random forest classification of the three cell types; (e) top 15 markers for classification obtained using Gini importance; (f), (g) and (h) density contour plots of the top 5 identified features taking two at a time, showing shifted density clusters for all the three cell types.

Next, a supervised learning approach using a random forest classifier was employed with the cell type label as target. Due to the presence of hundreds of decision trees, random forest algorithm prevents overfitting and cancels out errors from individual trees leading to a better generalized model. Our model showed high prediction accuracy (>90%) for all three cell types. The confusion matrix depicted in figure 3(d) shows our model’s strong accuracy: 98.87%, 93.46% and 90.28% in predicting neutrophils, PBMCs and RBCs respectively. A cluster map of the top 20 features for identified three clusters is shown in Figure S4(d). Again, Gini importance was utilised to find the top 15 markers used by the random forest classifier to classify the three cell types, which is demonstrated in figure 3(e). T_e_, Y Amp BC+ 2.3 MHz and T_en_ were found to be the top three markers contributing the most to the classification. Figure 3(f), (g) and (h) present the density contour plots of T_e_ vs Y Amp BC+ 2.3 MHz, T_en_ vs Y Amp BC+ 2.3 MHz and X Amp AC 8 MHz vs Y Amp BC+ 2.3 MHz respectively, which clearly show shifted contours for neutrophils, PBMCs and RBCs.

After identifying the top parameters for classification from the machine learning model, individual plots of the parameters are presented as follows. Figure 4(a) and (b) depict the histogram and the box plot of the reactive impedance amplitude obtained at BC+ electrode at 2.3 MHz, Y Amp BC+ 2.3 MHz. The histogram shows three distinct peaks for the three cell types, i.e., neutrophils, PBMCs and RBCs. Both Y Amp BC+ and Y Amp BC-at 2.3 MHz show similar trend in identifying distinct distributions for all the three cell types, and one of those amplitudes, i.e., Y Amp BC+ 2.3 MHz has been presented here. Figure S6 shows the histogram for Y Amp BC-2.3 MHz. Hence, Y Amp BC at 2.3 MHz stands out as one good distinguishing marker. From biological perspective, the reactive impedance at 2.3 MHz belongs to the β-dispersion regime which is strongly influenced by the cell membrane capacitance. Neutrophils are granular and have a highly folded membrane, which increases the surface area of the cell membrane resulting in higher effective capacitance. PBMCs and RBCs have smoother membranes which results in smaller membrane capacitance values than neutrophils.

**Figure 4:**
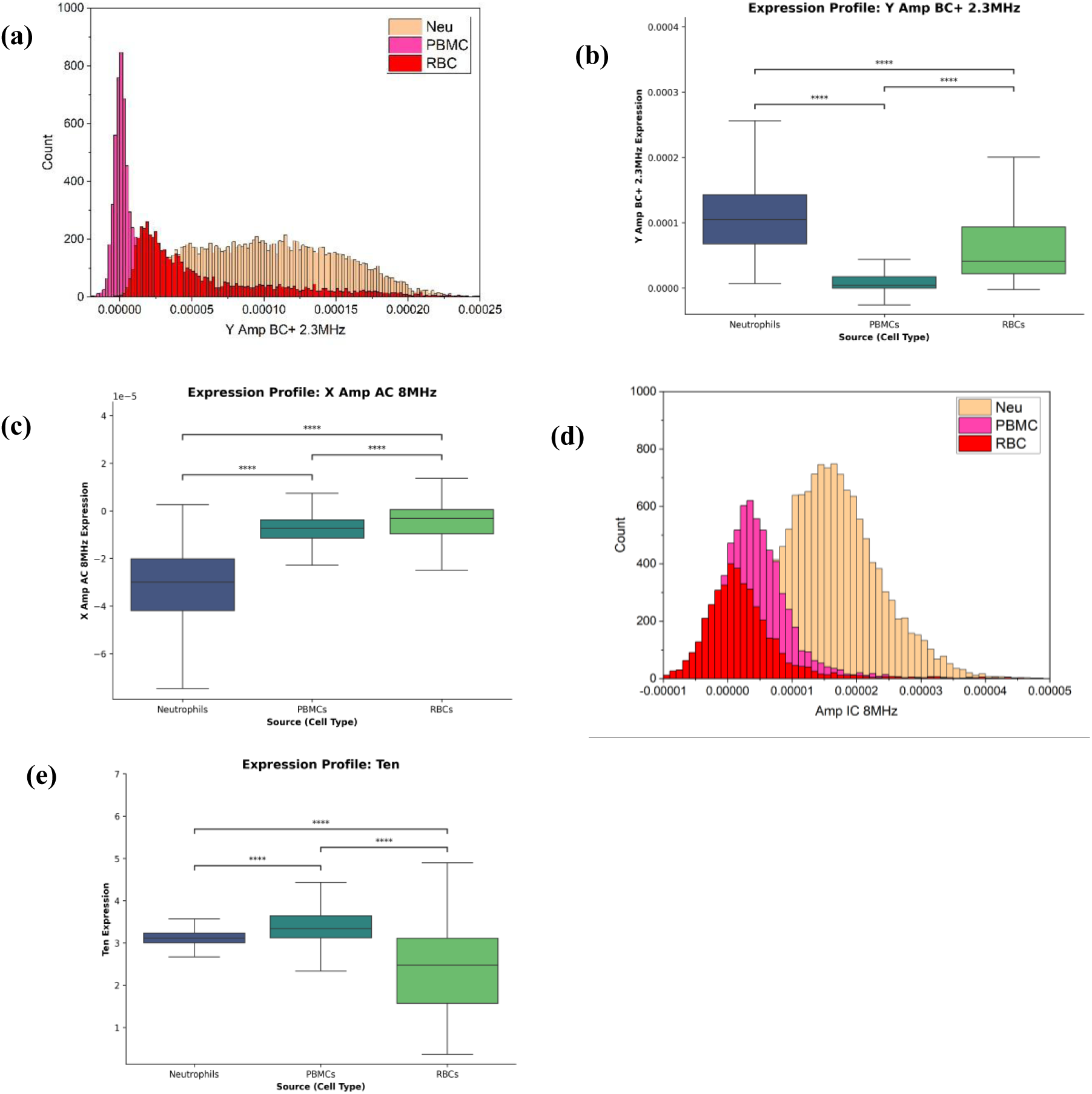
Experimental results from Neutrophils, PBMCs and RBCs (a) histogram of Y Amp BC+ at 2.3 MHz for neutrophils, PBMCs and RBCs; (b) Box plot for Y Amp BC+ 2.3 MHz with statistical significance by p value; (c) Box plot of X Amp AC 8 MHz with statistical significance; (d) histogram of Amp IC 8 MHz; (e) Box plot of entry time, Ten with statistical significance.

Figure 4(c) shows the box plot of the real part of the complex impedance obtained at AC electrode at 8 MHz, X Amp AC 8 MHz. The p<<< 1 validates it as another important marker for cell differentiation. Figure 4(d) is a histogram of real part of impedance at IC electrode at 8 MHz, Amp IC 8 MHz, which shows good distinction between neutrophils and PBMCs, but shows a significant overlap between PBMCs and RBCs, highlighting the marker as important for the former two cell differentiation but not for the latter two. According to the dielectric dispersion regime^[25]^, frequencies ranging from 7-20 MHz probe the cytoplasm. We chose 8 MHz as the working frequency because it experimentally showed the highest SNR in that range. RBCs have a high density of mobile ions in their cytoplasm, leading to the highest cytoplasmic conductivity, and hence, lowest resistivity, leading to the lowest amplitude at 8 MHz. PBMCs contain a large nucleus which is less conductive than the surrounding cytoplasm, leading to lower conductivity than RBCs, and hence, higher amplitude. Neutrophils are packed with granules having individual membranes. These membranes act as capacitive barriers and lower the overall conductivity of the ions inside the cytoplasm.

Figure 4(e) depicts the box plot for the entry time, T_en_, for the three cell types, again establishing as an important distinguishing parameter with p<<<1.

### 3.2 Co-flow of Neutrophils and PBMCs

After verifying the individual flow of cells, a mix population sample was put to test. Neutrophils and PBMCs were extracted separately from the density centrifugation method and then mixed in 1:1 volume ratio in PBS buffer to a final concentration of 1.5×10^3^ cells/µL.

#### 3.2.1 Direct peak analysis

Fig 5(a) and 5(b) show histograms of Y Amp BC+ and Y Amp IC for PBMCs and neutrophils at 2.3 MHz from the individual flow experiment. Both the cell types exhibit distinct peaks, with PBMCs extending a tail-like behaviour probably corresponding to monocytes, and PBMCs possess a lower peak amplitude than neutrophils. Figure 5(c) shows the box plot for the mean value of Y Amp IC at 2.3 MHz with distinct mean and standard deviation for both neutrophils and PBMCs. Figure 5(d) depicts the error bar plot of Max Amp BC^1/3^ with distinct mean and standard deviation for both the cell types.

**Figure 5:**
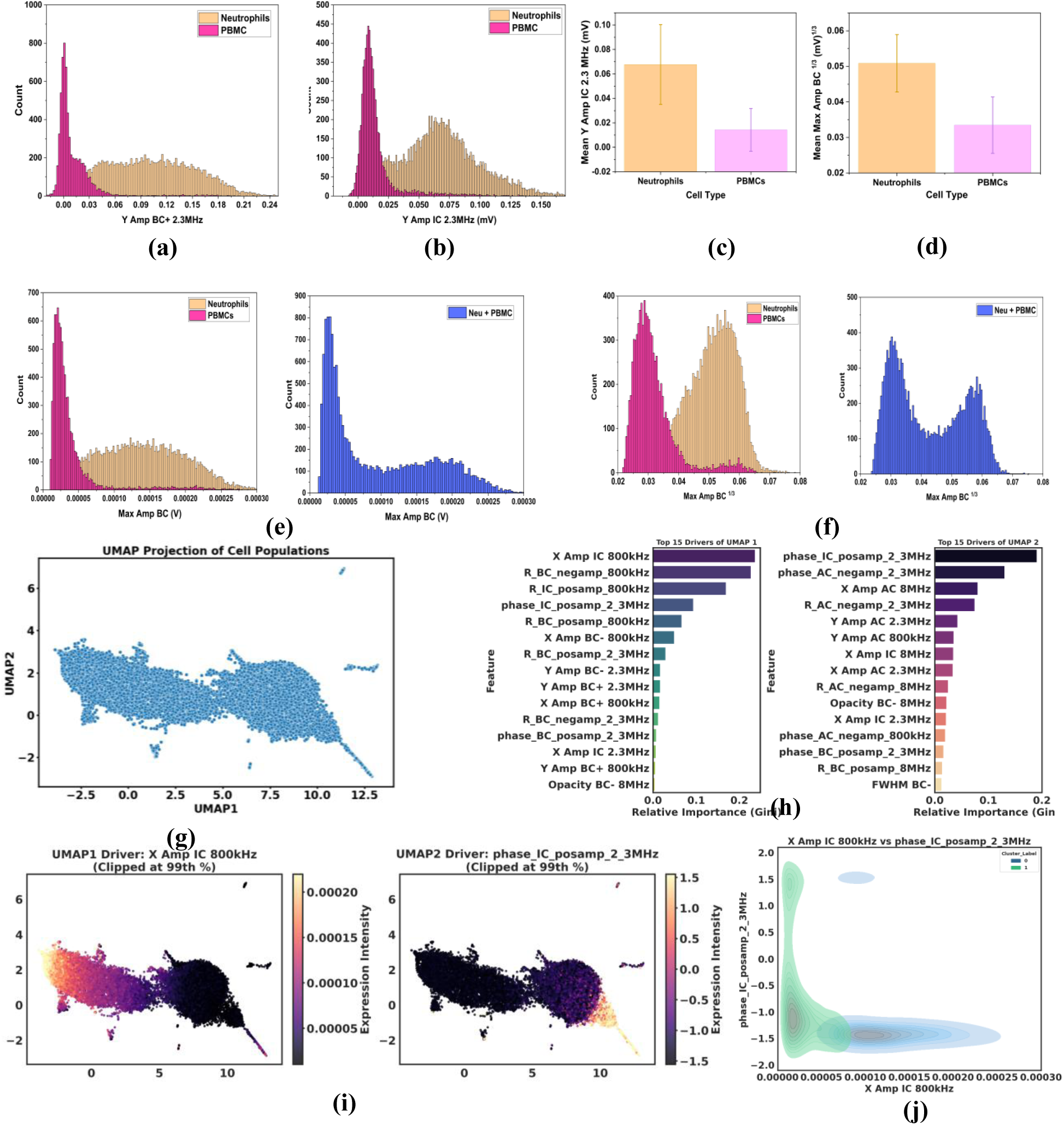
Experimental and ML analysis results for co-ffow of neutrophils and PBMCs: (a) and (b) histograms of two parameters, Y Amp BC+ and Y Amp IC at 2.3 MHz respectively, depicting two distinct distributions; (c) and (d) box plots for Mean Y Amp IC 2.3 MHz and Mean Max Amp BC^1/3^ at 800 kHz, showing two separate populations; (e) and (f) histogram comparison of Max Amp BC and Max Amp BC^1/3^ for (left) individual ffow of neutrophils and PBMCs and (right) combined ffow of the two cell types with the right plot showing the same peak values; (g) UMAP projection scatter plot of co-ffow depicting two clusters validating the presence of two populations; (h) top 15 drivers of each of the UMAP coordinates obtained using Gini importance; (i) overlap of the top features of UMAP1 and UMAP2 on the UMAP projection plot; (j) density contour plot of the top features of each of the UMAP showing two distinct clusters.

Next, we compared the distributions of individual cell flow with the mixed population flow. Fig 5(e) (left) shows the histogram of Max Amp BC for the two cell types flown separately. In comparison, Fig 5(e) (right) shows the histogram of Max Amp BC for the mixed population. The mixed population histogram shows two gaussian peaks with the same peak values as the individual cell populations, 0.021 mV for PBMCs in separate vs 0.025 mV in mixed and 0.14 mV for neutrophils in separate vs 0.184 mV in mixed, thereby confirming the presence of PBMCs and neutrophils. Similarly, fig 5(f) shows the histograms of the Max Amp BC^1/3^ for the two cell types, both individually flown and in mixed population. Again, the peak values for both the cell types are similar for both the flow scenarios: 0.028 mV^1/3^ in separate vs 0.03 mV^1/3^ in mixed population for PBMCs and 0.055 mV^1/3^ in separate vs 0.05 mV^1/3^ in mixed population for neutrophils. However, the distribution for the Max Amp BC^1/3^ has an equivalent height-to-width ratio for the two cell types than Max Amp BC, and hence, is a better metric to classify them.

#### 3.2.2 Machine Learning analysis

After the preliminary peak data analysis, we implemented UMAP on the mix population peak data. The first scatter plot obtained for the 2D UMAP projection is shown in figure 5(g). Two separate clusters clearly indicate the presence of two different cell populations, thereby validating our experiment. Upon finding out the top 15 important features contributing to the UMAP coordinates, figure 5(h) presented real impedance magnitude at IC electrode at 800 kHz (X Amp IC 800kHz) and impedance phase at IC electrode at 2.3 MHz (phase_IC_posamp_2_3MHz) as the top UMAP drivers with the highest Gini importance levels. The impact of the top drivers on the UMAP projection scatter plot is depicted in figure 5(i). To illustrate the separation of clusters using the top UMAP drivers, figure 5(j) shows the density contour plot of X Amp IC 800kHz vs phase_IC_posamp_2_3MHz, which highlights two well separated clusters.

### 3.3 Flow of PBMCs

PBMCs were extracted from whole blood using density gradient centrifugation method, as described in previous subsections and diluted in 1x PBS to the final concentration of 1.5×10^3^ cells/µL.

#### 3.3.1 Peak analysis and machine learning analysis

Fig 6(a) shows the histogram of Opacity BC+ 2.3 MHz, calculated as the ratio of amplitude at BC+ electrode at 2.3 MHz to the amplitude at 800 kHz. Presence of two distinct peaks indicate two sub-populations present in PBMCs. We conducted a UMAP analysis of the PBMCs features whose density spread is shown in figure 6(b). Two bright zones validated our previous observation of two subpopulations. Furthermore, through the random regressor model whose bar graph shown in figure 6(c), Opacity BC+ 2.3 MHz was highlighted as the topmost feature contributing to the UMAP coordinates, marking it as the single most important feature for PBMC sub-classification. Post UMAP, we utilised an unsupervised clustering algorithm, hierarchical density-based spatial clustering of applications with noise (HDBSCAN), to identify clusters automatically by setting minimum cluster size to 1000. Figure 6(d) confirmed two primary subpopulations inside PBMCs with a potential third population which could either be a minority subpopulation or noise. We also created a heatmap of the top 10 features standardised through a z-score algorithm, depicted in figure 6(e).

**Figure 6:**
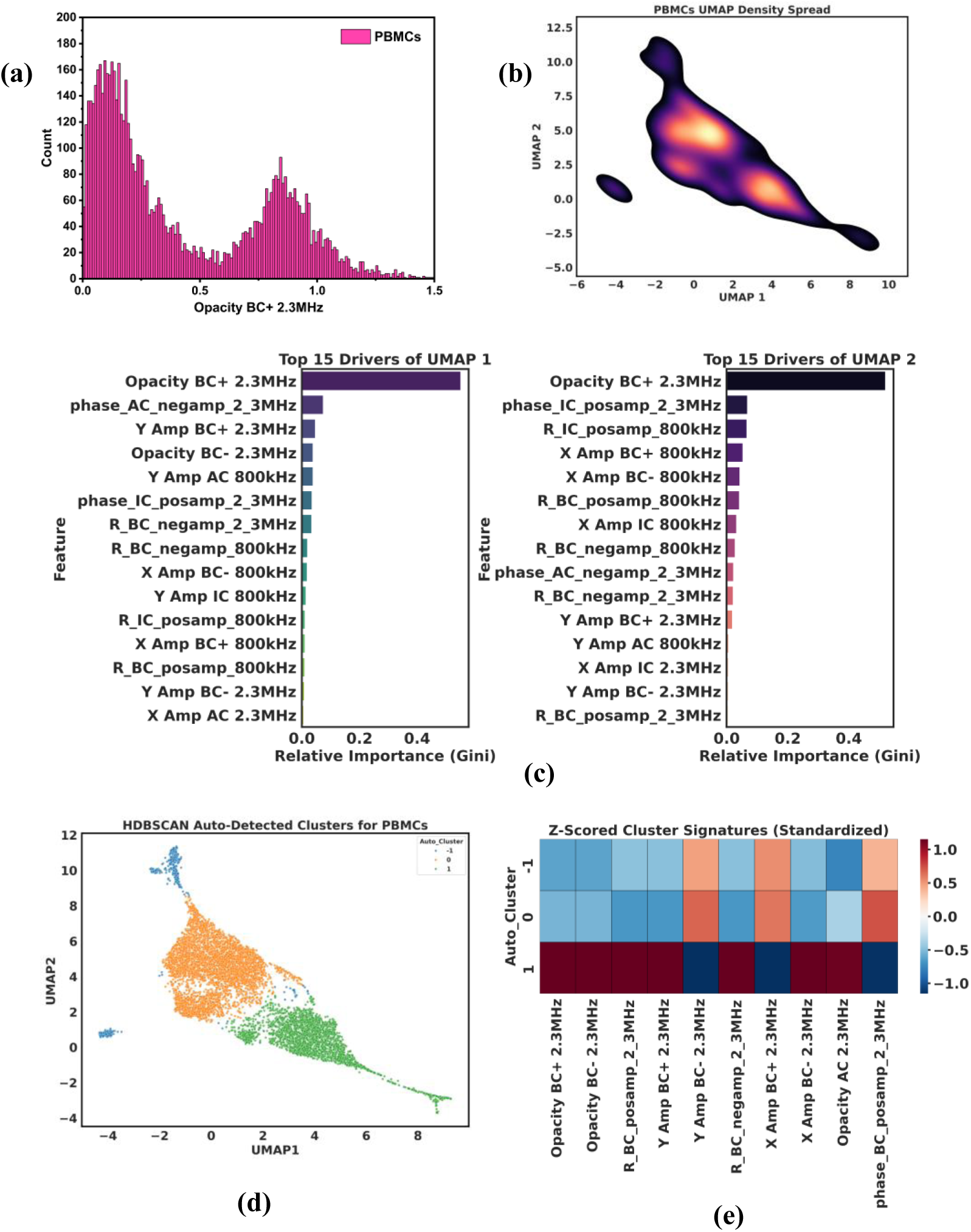
Experimental and ML analysis results for sub-populations of PBMCs (a) histogram for Opacity BC+ at 2.3 MHz showing two distinct peaks indicating the presence of at least two major sub populations; (b) UMAP density map showing two cluster centers validating the presence of two sub populations; (c) top 15 drivers of the UMAP coordinates, with the same top feature, highlighting the importance of Opacity BC+ 2.3 MHz alone as a powerful differentiating marker; (d) unsupervised clustering algorithm, HDBSCAN used to identify the presence of clusters in the UMAP scatter plot; (e) cluster map of the top 10 features standardized using z-scores.

Based on the model developed for the three cell types, namely neutrophils, PBMCs and RBCs, Figure 7 shows the workflow for a new sample. The sample comprising of those three cell types flowed through the designed microfluidic device and analysed through our trained model will successfully classify the cells using both their electrical and mechanical properties. The top 5 classification parameters identified by our model are Y Amp BC 2.3 MHz, Exit time, Opacity BC 2.3 MHz, Entry time, and X Amp AC 8 MHz.

**Figure 7:**
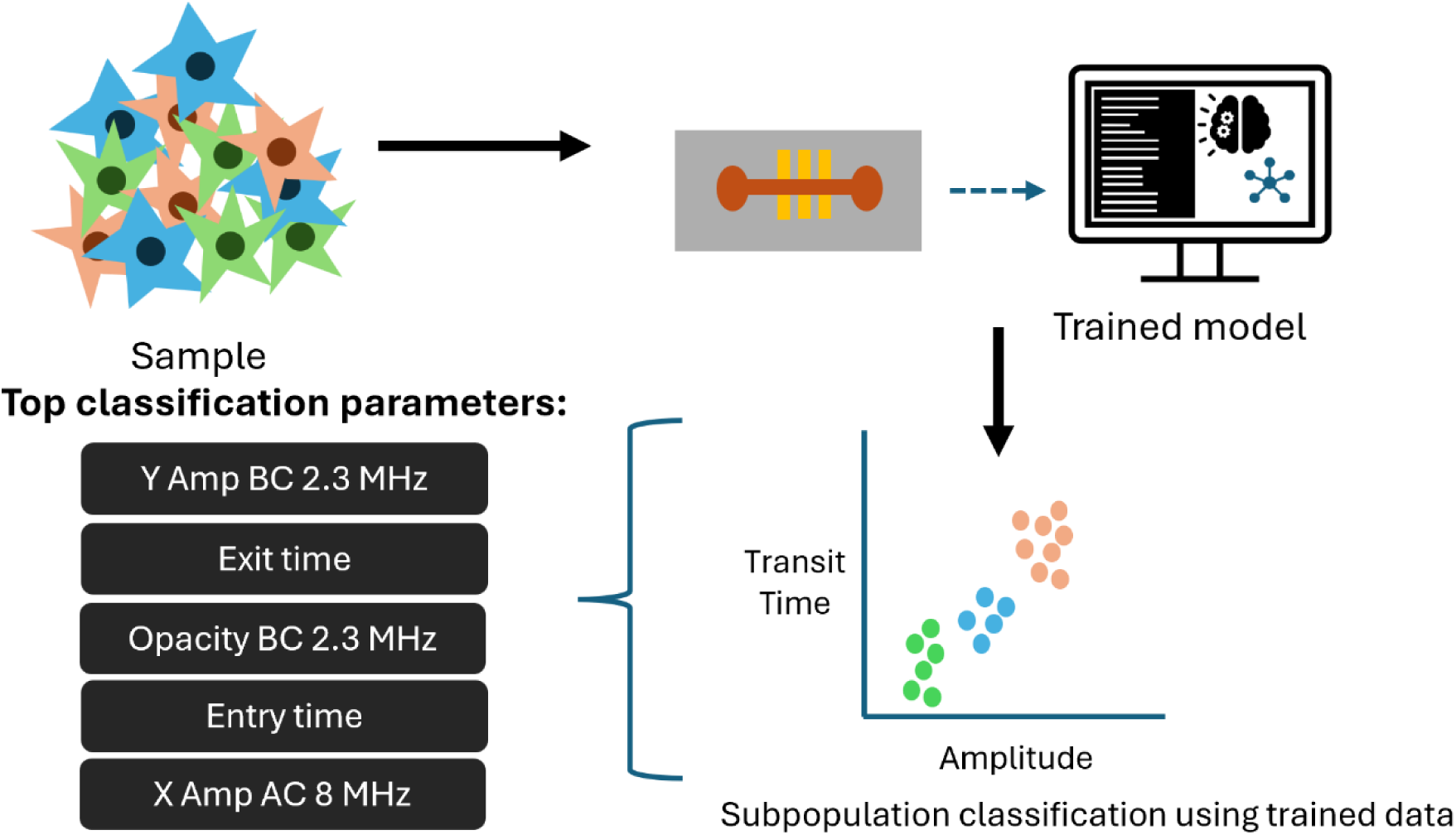
Workflow for a new cell sample.

## 4. Conclusion

In this study, we demonstrated a multi-frequency mechanical and impedance cytometer capable of understanding internal cell components at multiple stages of deformation: native, deformed and post-deformed cellular states. Our design is utilised to distinguish between multiple blood cell types, integrated with unsupervised and supervised machine learning models, with up to 98% classification accuracy. Our platform is also capable of identifying multiple cell types within a mixed population with strong correlation with the independent cell data. In our experiments, we obtained a high throughput of 3.8k cells/min. In conclusion, the multi-frequency electro-mechanical cytometer leverages cellular impedance of various intracellular components during different stages of deformation to distinguish between multiple cell types.

## 5. Methods

### Cell preparation

Blood samples were collected in ethylenediaminetetraacetic acid (EDTA) coated vacutainer tubes from healthy volunteers. The leukocytes, namely, neutrophils and peripheral blood mononuclear cells (PBMCs), were extracted from whole blood using the standard density gradient centrifugation method (Figure S1). Briefly, the process includes layering blood onto Histopaque-1077 carefully in a centrifuge tube, ensuring no mixing, and centrifuging it for 30 minutes with no brake condition. The centrifuge process yields four separate layers: bottom layer with red blood cells (RBCs) and neutrophils, first middle layer with Histopaque-1077, second middle layer with PBMCs, and the topmost layer with plasma. PBMCs are carefully pipetted out and diluted in phosphate buffered saline (PBS). The mixture of neutrophils and RBCs is treated with RBC lysis buffer, followed by neutrophils separation through centrifugation, and dilution in PBS.

### Device Design and fabrication: Microfluidic device

The microfluidic device was created using a master mould by soft lithography process, where the master mould was fabricated using photolithography. Figure S2 shows the process flow to fabricate the master mould followed by the PDMS replica. To create the master mould, a clean silicon wafer was spin-coated with a positive photoresist and patterned using optical lithography. Later, the patterns were etched using deep reactive ion etching and the required depth was verified using stylus profilometer. After obtaining the master mould, a negative replica of the patterns was created by pouring poly di-methyl siloxane (PDMS) onto the wafer and cured.

The dimensions of the channel were kept 20 µm (width) x 15 µm (depth) and the constriction dimensions were 7 µm (width) x 15 µm (depth).

### Microelectrode fabrication

The microelectrodes were fabricated on a glass slide using photolithography and deposition. Figure S3 shows the process flow for microelectrodes fabrication. Glass slide was cleaned using piranha solution, comprising of concentrated sulfuric acid, H_2_SO_4_ and 30% hydrogen peroxide, H_2_O_2_, in the ratio 3:1 to remove organic and metallic contaminants. Next, a positive photoresist was coated and patterned using optical lithography, followed by deposition of metal layers of Cr/Au of thicknesses 20 nm/100 nm using e-beam evaporation. Eventually, excess metal was removed using lift-off and microelectrodes were obtained. The dimensions of the electrodes were kept 15 µm width with 15 µm gap.

### PDMS bonding

The microelectrodes and microfluidic device were bonded together by plasma treatment after carefully aligning under the microscope. To amplify the cell impedance signals, the contact pads of the electrodes were epoxy-bonded and soldered to an adaptor board which connects to an in-house developed instrumentation amplifier board.

### Experimental setup and data acquisition

The setup consisted of a microfluidic device, instrumentation amplifier board, a lock-in amplifier, and a computer. The packaged microfluidic device sat on top of the instrumentation amplifier board. The in-house developed instrumentation amplifier was a two-channel low-noise amplifier utilising an operation amplifier (AD8066) by Analog Devices. A lock-in amplifier, HF2LI by Zurich Instruments was used to remove noise and perform multi-frequency measurement and was controlled using LabOne software by Zurich Instruments. Each passing cell was probed at three frequencies (800 kHz, 2.3 MHz and 8 MHz) simultaneously. The data was sampled at 14k samples per second, and the low-pass filter bandwidth was set to 1 kHz. The resulting data was processed using a custom written Python script to identify cell transit events automatically.

## Ethical Declaration

Permission to work with human blood was obtained from the Indian Institute of Science institute health ethics committee (IHEC) after submitting detailed information about the experimental procedure. All blood samples were collected from healthy volunteers after getting informed consent forms signed by them.

## Acknowledgements

We acknowledge funding support from Program for Research and Application in Micro and Nanotechnologies (PRAMAN) scheme by the Ministry of Electronics and Information Technology (MeitY), Government of India (Grant Number: SP-MEIT-24-0007). IB acknowledges Prime Minister’s Research Fellowship (PMRF), Ministry of Education (MoE) for providing financial support. We thank Dr. Ramray Bhat and Dr. Siddharth Jhunjhunwala from Department of Bioengineering, IISc for allowing us to use their cell culture facility for blood cell separation. We thank Dr. Aditya Malladi from the Health Centre, Indian Institute of Science (IISc) and lab technicians for their help in obtaining blood samples. We thank the facility technologists at national nano fabrication centre (NNFC) and packaging and systems facility (PASF) for their valuable inputs and help in fabrication and device assembly.

