## Supplementary info for "Machine learning assisted multi-level differentiation of blood cells using multi-frequency electro-mechanical impedance cytometry"

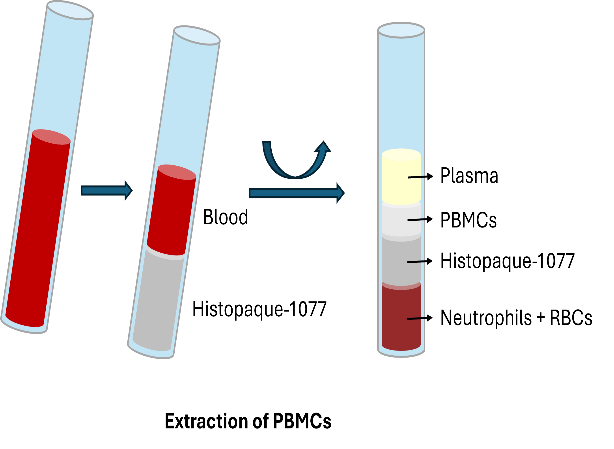

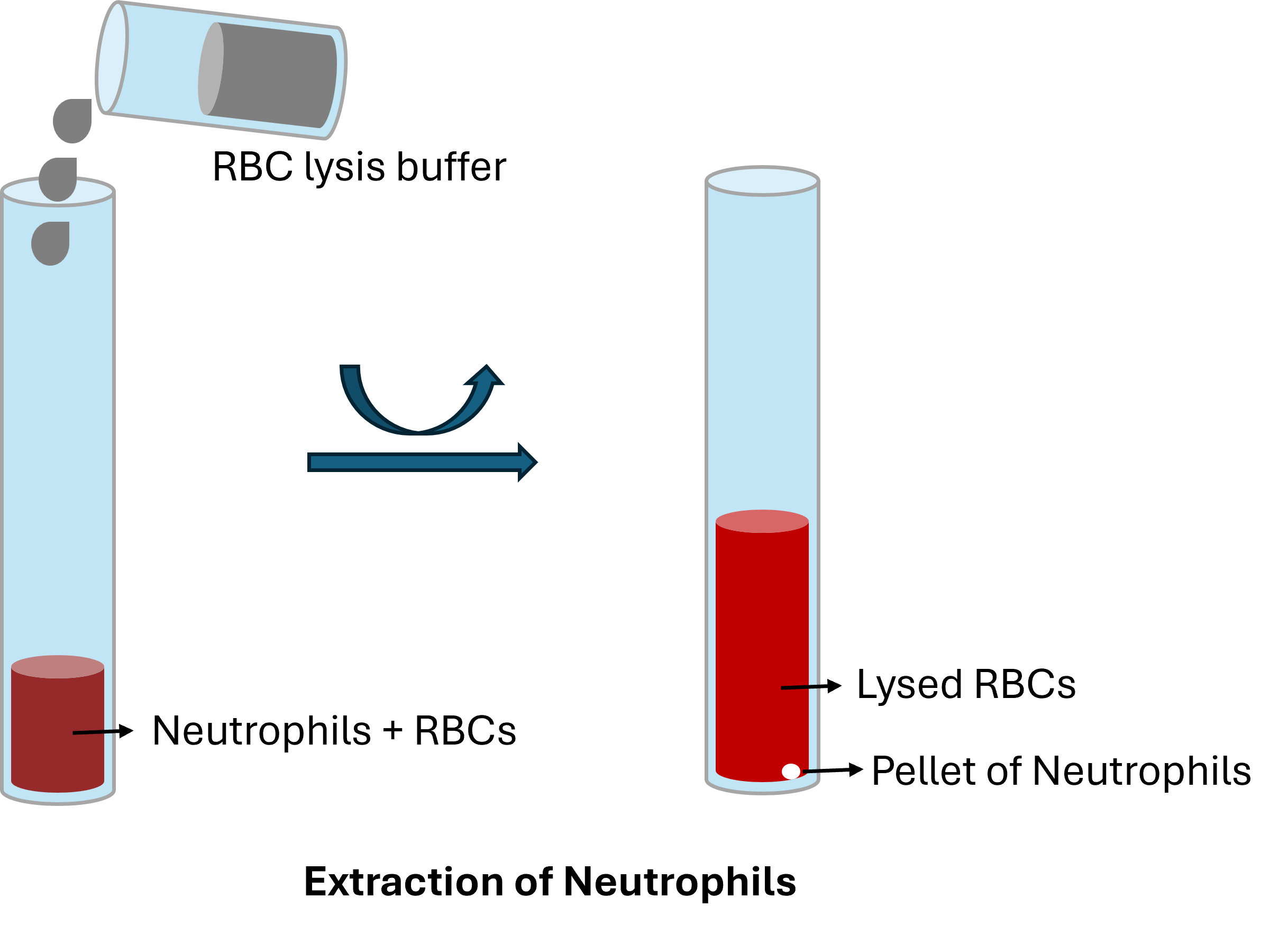

*Figure S1: Cell extraction process flow using Histopaque-1077. Blood layered onto Histopaque and centrifuged to obtain a PBMC layer, leaving neutrophils and RBCs sedimented. Further, RBCs are lysed from the mixture to obtain neutrophils. Later, the cells are diluted in PBS buffer.*

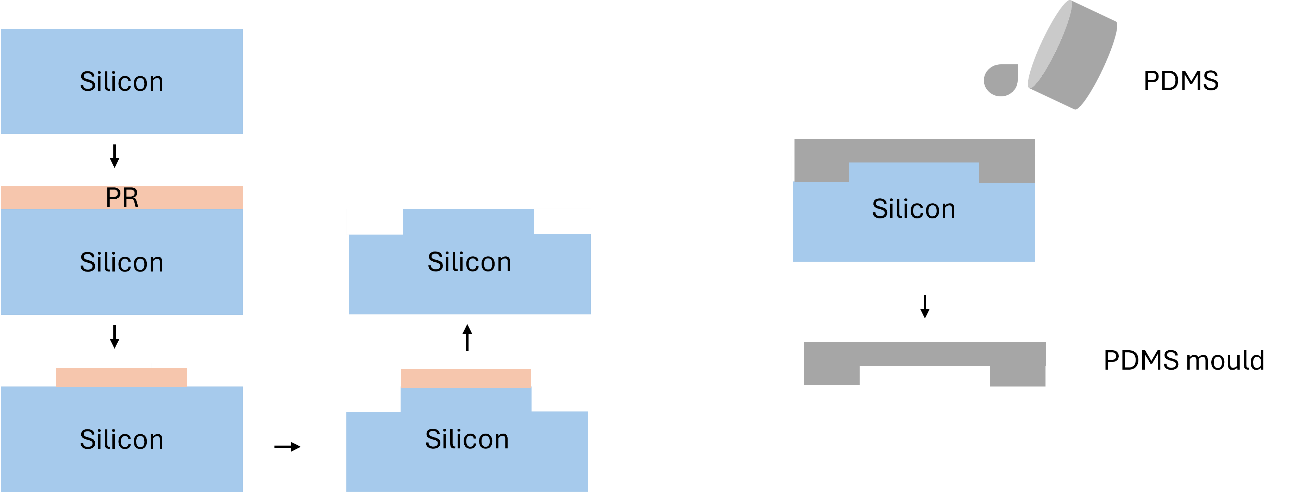

*Figure S2: Process flow for microfluidic device fabrication*

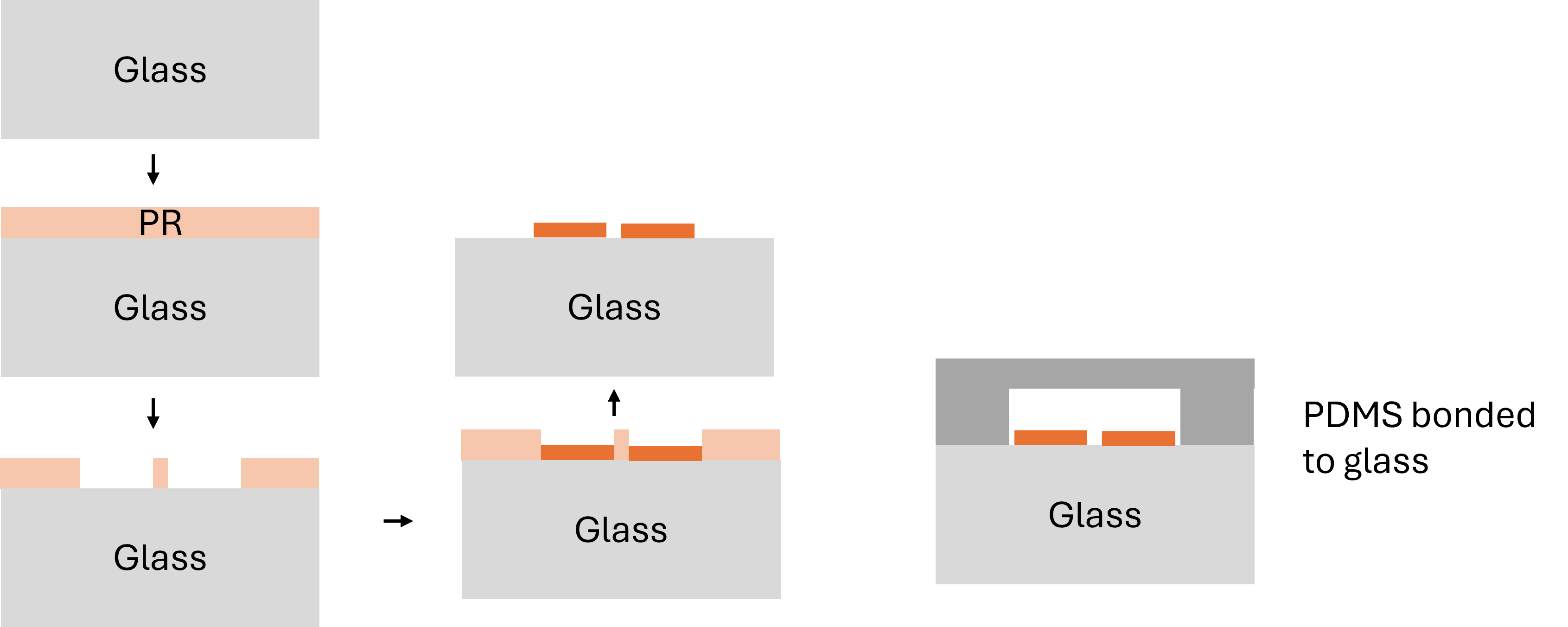

*Figure S3: Process flow for microelectrode fabrication*

**Machine Learning Workflow:**

Figure S4 shows the machine learning workflow for cell data analysis. From preliminary data analysis and peak detection, 72 features/cell were extracted. A cluster map of the 72 features is shown in figure S4(a). These were fed as input to a dimensionality reduction technique, UMAP, to visualize clusters present in the data, whose two coordinates: UMAP1 and UMAP2 are plotted in figure S4(b). After verification of clusters, HDBSCAN clustering algorithm was used to identify clusters and a random forest regressor was trained on the cluster identity target label and top 5 features contributing to the clusters were identified (figure S4(c)). After the verification of clusters, a random forest classifier was used with train test split of 80:20. The classifier found the top 20 features essential for classification which are shown in figure S4(d).

*Figure S4: Illustration of the machine learning workflow developed for cell classification: The first figure shows the cluster map with the original 72 features for 100 cells, showing huge variation in features across cells. Next, dimensionality reduction converts 72 features to 2 UMAPs to visualise the presence of clusters in the data. Top 5 features which drive the clustering are shown for representation in the third figure. Lastly, a random forest classifier is trained and tested on the data and a cluster map of the top 20 features identified by the classifier are presented.*

**(b)**

**(d)**

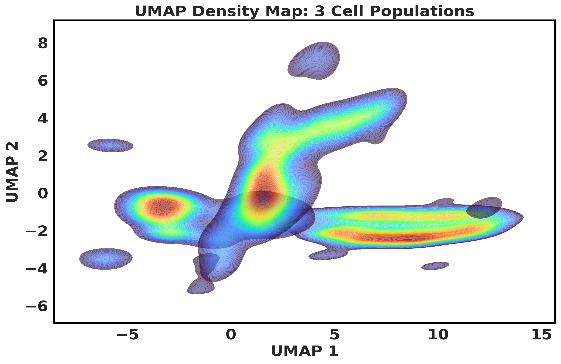

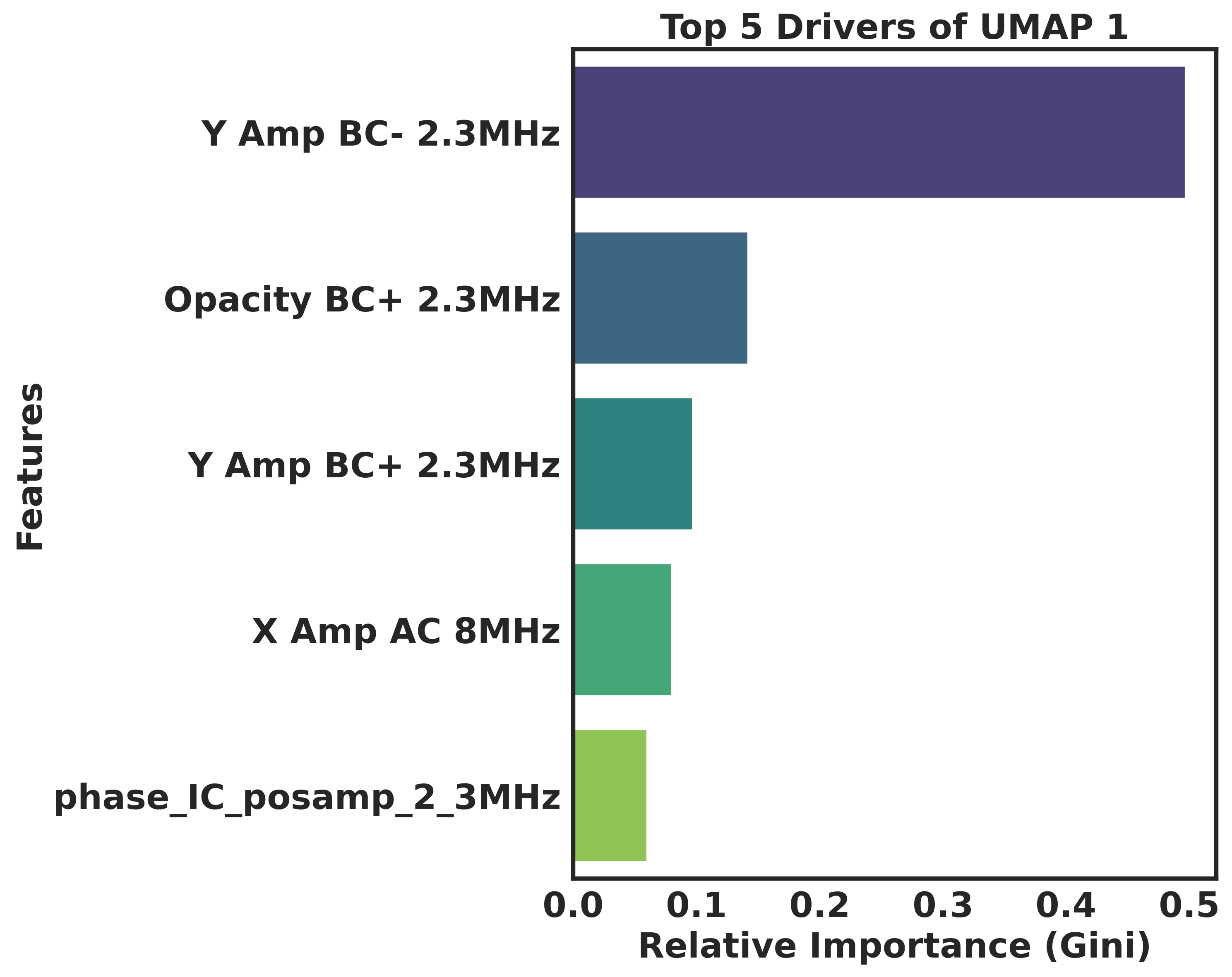

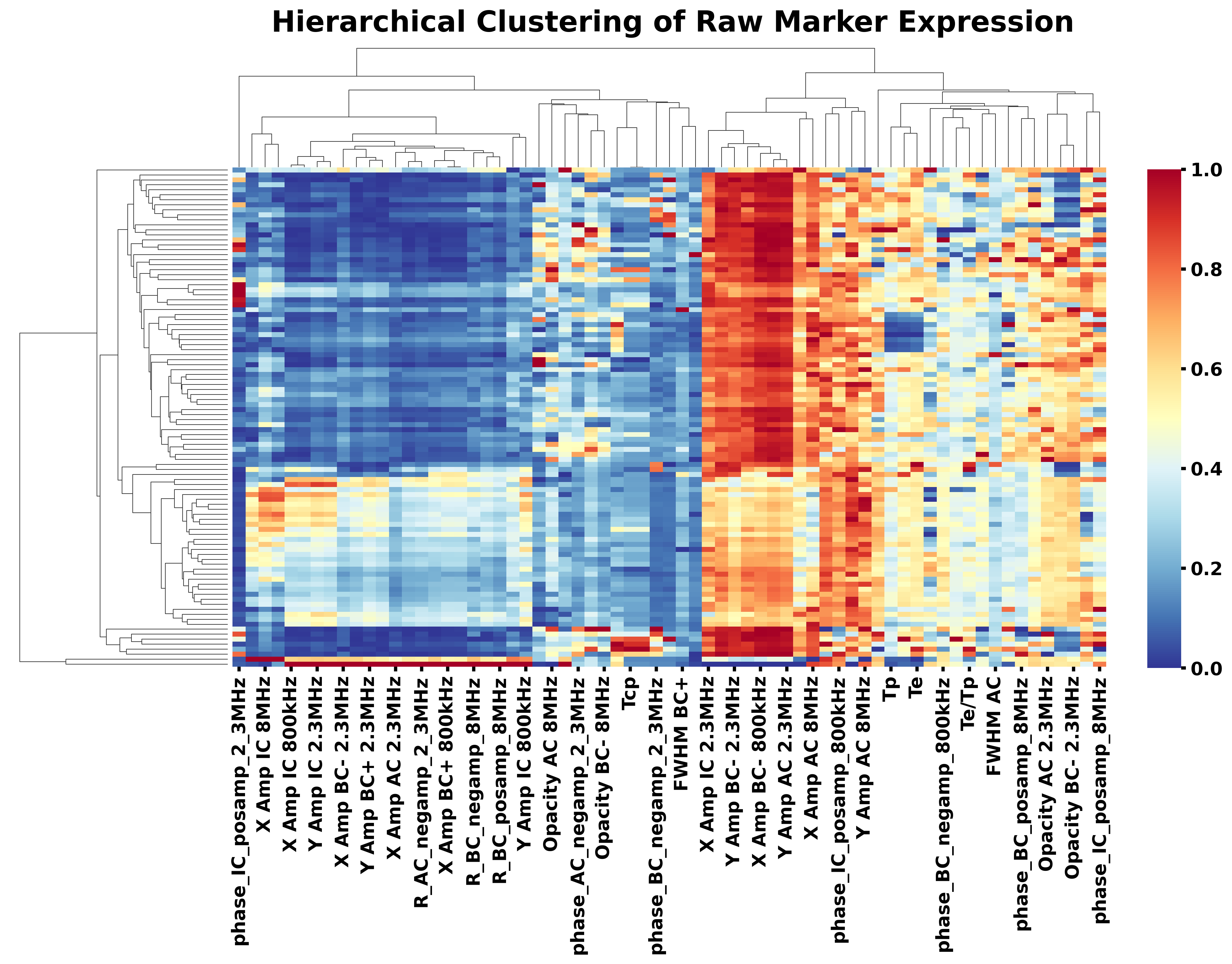

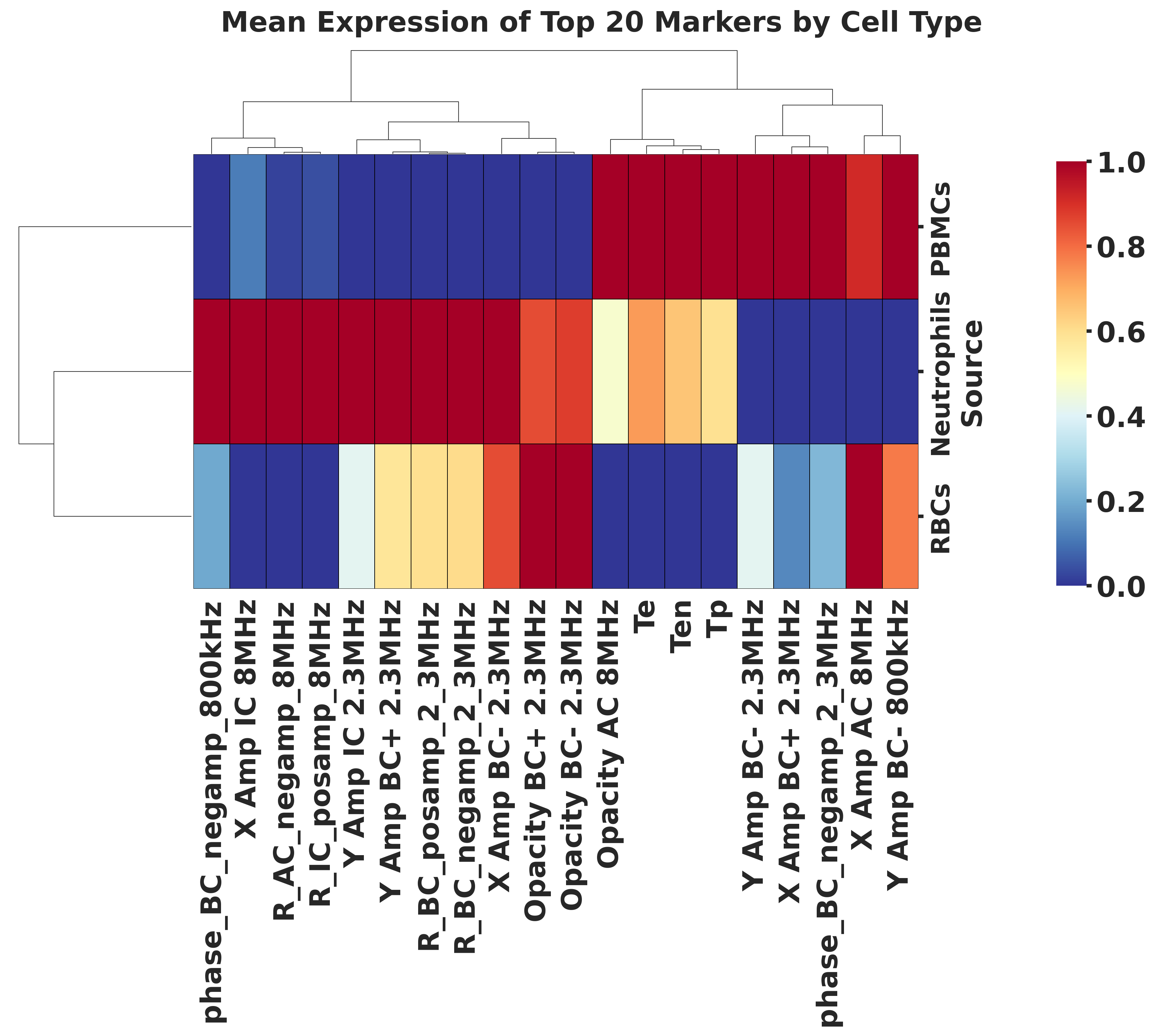

Cluster map of 72 input features

UMAP density plot depicting multiple clusters present in the data

Top 5 features driving the clustering

Cluster map of top 20 features responsible for classification

**(c)**

**(a)**

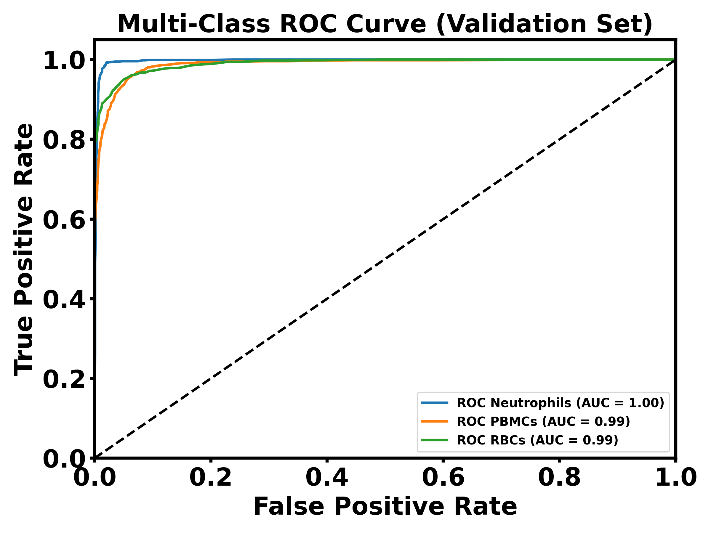

*Figure S5: AUC curve for flow of neutrophils, PBMCs and RBCs showing high precision and recall, AUC ~ 1*

*Figure S6: Histogram of Y Amp BC- 2.3 MHz for neutrophils, PBMCs and RBCs showing three different distributions*

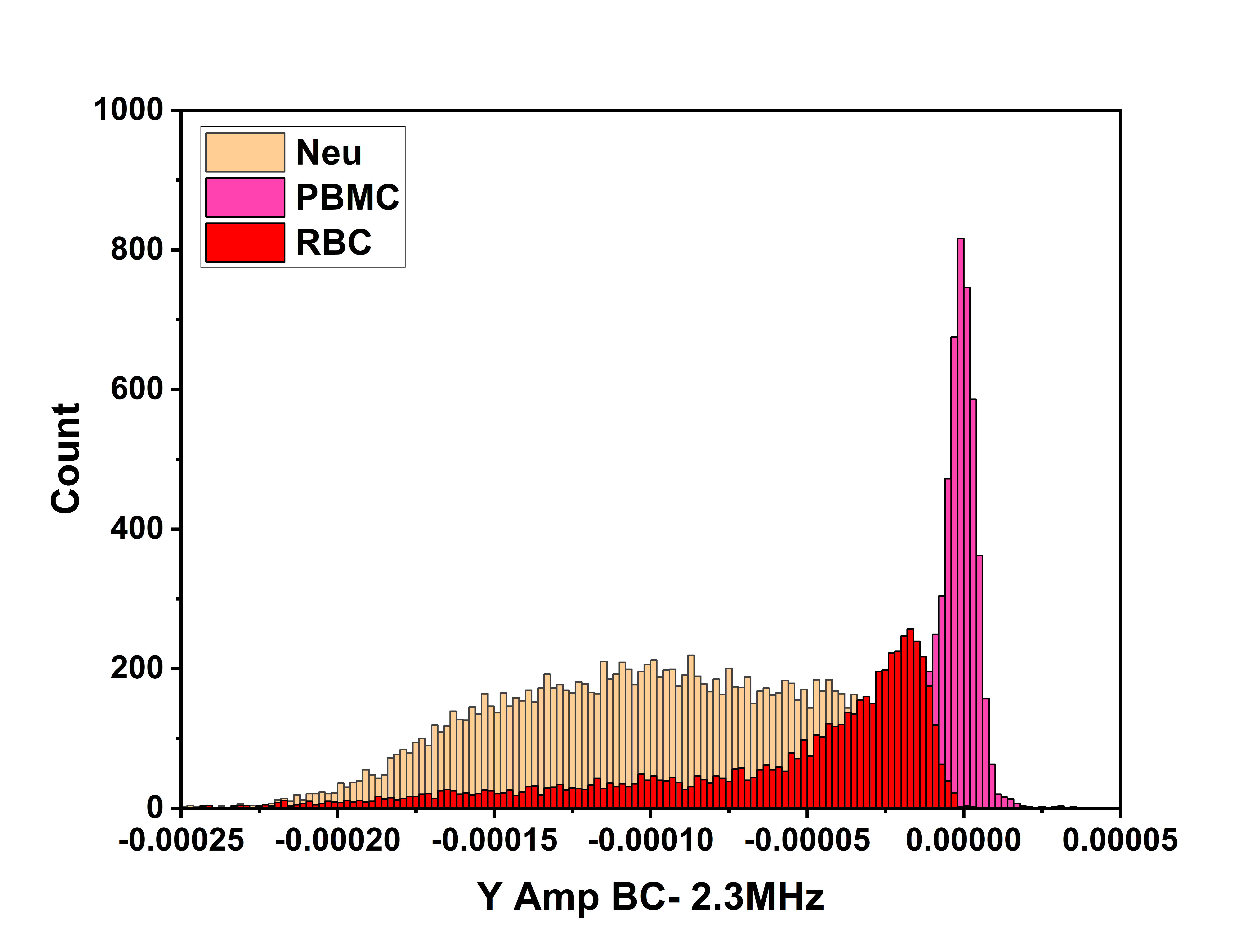

**Random Forest Hyperparameter tuning:**

To maximise the predictive power of random forest classifier, we performed hyperparameter tuning using a randomised search algorithm (RandomizedSearchCV) with 3-fold cross validation. The following parameters in random forest were optimized:

1. *n_estimators* (100, 500): The **number of trees in the forest** were explored to reduce variance without increasing computational overload.
2. max_depth ((10, 50) or None): The **maximum depth of the tree** was varied once in the range (10,50) and the other time with no limit. By testing both shallow and deep trees, the model captures complex patterns without overfitting.
3. *min_samples_leaf* (1,20): The **minimum number of samples present in a leaf node** were explored in the range of (1,20) to prevent overfitting and applying rules which are statistically significant.
4. *max_features* (‘sqrt’, ‘log2’, None): The **number of random subset of features** to be considered at each split were varied to prevent any feature to dominate over the others.
5. *class_weight* (‘balanced’, ‘balanced_subsample’): Addresses **data imbalance** if certain cell type is rarer than the other.

We ran 25 iterations of the above optimisation and found the following hyperparameters to have the highest mean cross-validation accuracy:

*n_estimators* = 317

*max_depth* = 22

*min_samples_leaf* = 1

*max_features* = None

*class_weight* = ‘balanced’

Table 1: Description of electrical and mechanical parameters obtained through the system.

| **Parameter acronym** | **Parameter description** |
| --- | --- |
| Te | Exit time (Time to exit the constriction) |
| Ten | Entry time (Time to enter the constriction) |
| Tp | Passage time (peak to peak time before constriction) |
| Y Amp BC+/BC- 2.3 MHz  Generic form:  <Type of Impedance> Amp <electrode set> <frequency>  where:  Type of impedance ⊂ {Resistive (X), Reactive (Y)}  electrode set ⊂ {BC+, BC-, IC, AC}  Frequency ⊂ {800 kHz, 2.3 MHz, 8 MHz} | Reactive impedance amplitude measured at one of the before constriction electrodes at 2.3 MHz |
| X Amp AC 8 MHz  Generic form:  <Resistive impedance amplitude> Amp <electrode set> <frequency> | Resistive impedance amplitude measured at after constriction electrode at 8 MHz |
| Opacity AC 8 MHz  Generic form:  Opacity <electrode set> <frequency> | Opacity (Ratio of net amplitude at high frequency to amplitude at 800 kHz) measured at after constriction electrode at 8 MHz |
| R_BC_posamp_2_3MHz  Generic form:  R_<electrode set>_<amplitude polarity>_<frequency>  where:  amplitude polarity ⊂{posamp (BC+, IC), negamp (BC-, AC)} | Net impedance amplitude  ($R= \sqrt[2]{X^{2}+ Y^{2}}$) measured at the first before constriction electrode (BC+) at 2.3 MHz |
| phase_IC_posamp_2_3MHz  Generic form:  phase_<electrode set>_<amplitude polarity>_<frequency> | Phase of impedance ($phase= \tan^{-1} \frac{Y}{X}$) measured inside constriction electrode at 2.3 MHz |
